# Use of steric blocking antisense oligonucleotides for the targeted inhibition of junction containing precursor microRNAs

**DOI:** 10.1101/2024.04.08.588531

**Authors:** Sicong Ma, Samantha A. Howden, Sarah C. Keane

## Abstract

Antisense oligonucleotides (ASOs) are widely used as therapeutics for neurodegenerative diseases, cancers, and virus infections. One class of ASOs functions to enhance protein expression by sequestering the mature microRNA (miRNA) in a double-stranded structure within the RNA-induced silencing complex (RISC). An alternative approach for the targeted control of gene expression is to use ASOs that bind to the pre-elements of miRNAs (pre-miRNAs) and modulate their enzymatic processing. Here, we demonstrate that ASOs can be used to disrupt a specific structural feature, “junction,” within pre-miR-31 that is important in directing efficient processing by the Dicer/TRBP complex. Furthermore, we extend and validate this strategy to pre-miR-144, which has a similar junction-dependent structure-function relationship. We found that a significant number of human pre-miRNAs are predicted to contain junctions, and validated our ASO approach on several members of this group. Importantly, we also verified the application of junction-targeting ASOs for the specific inhibition of pre-miRNA processing *in cell*. Our study reemphasizes the important roles of RNA structure in regulating Dicer/TRBP processing of pre-miRNAs and provides the framework to develop structure-informed ASOs that serve to inhibit miRNA production.

## INTRODUCTION

Antisense oligonucleotides (ASOs) are a type of chemically modified, short, single-stranded nucleic acid that bind to target RNAs via Watson-Crick base pairing. ASOs are typically 12-30 nucleotides (nts) long and increasing the ASO length results in increased specificity for the target RNA. The mechanisms through which ASOs regulate RNA function can be broadly categorized as either promoting RNA turnover or sterically blocking the targeted RNA which can affect a multitude of processes (1–3).

One common use of ASOs is to promote target RNA cleavage by recruitment of either RNase H1 or Argonaute 2 (Ago2) proteins (4). RNase H1 is an endogenous endonuclease that is present in the nucleus, cytoplasm, and mitochondria of mammalian cells (5). RNase H1 plays as important role in mitochondrial genome replication and also functions in genome maintenance by removing R-loop (5,6). RNase H1 recognizes RNA-DNA heteroduplexes and catalyzes a phosphoryl transfer reaction that results in a break in the phosphodiester backbone of the RNA (7). Exogenous small interfering RNAs (siRNAs) can also be used for nucleic acid therapy (4). Similar to a mature microRNA duplex, siRNA duplexes, which may contain a variety of chemical modifications (8), can be loaded into Ago2 to form the RNA-induced silencing complex (RISC). The functional RISC binds to the 3ʹ-untranslated region (UTR) of a specific messenger RNA (mRNA) which induces cleavage of that mRNA (9,10).

Steric blocking is another approach for ASO-targeted inhibition of RNA function. Mammalian mRNAs undergo a number of post-transcriptional modifications, including the splicing of one or more intron sequences out of a precursor mRNA (11). Splicing is a fundamental process in RNA biology and the dysregulation of splicing is associated with numerous diseases and pathologies (12). ASOs can be used to target the splicing process, broadly regulating gene expression. Duchenne muscular dystrophy (DMD) is a severe neuromuscular disease that is caused by loss-of-function mutations in the dystrophin gene (13). Eteplirsen, a morpholino ASO, is designed to skip exon 51, which leads to the production of a partially functional dystrophin protein (14). Conversely, Nusinersen, an ASO containing modifications at both the 2’-hydroxyl and the phosphate backbone, functions to reduce exon skipping and has been shown to be an effective treatment of spinal muscular atrophy, a rare neuromuscular disorder (15). Nusinersen increases levels of survival of motor neuron 2 (SMN2) mRNAs that include exon 7 which can be translated into a functional SMN protein (15,16). Steric blocking ASOs that target RNA elements in the 5ʹ-UTR of mRNAs can function to up- or down-regulate protein translation. For example, ASOs that overlap with or are close to the mRNA start codon can prevent ribosome binding or assembly on the mRNA, reducing translation (4,17). Alternatively, ASOs that target repressive elements, like an upstream open reading frame (uORF) or stem-loop structure within the 5ʹ-UTR, can enhance the translation of the downstream gene (18,19).

Finally, ASOs can be used to sequester mature microRNAs (miRNAs) to enhance protein expression (1,4,20). In the nucleus, RNA polymerase II transcribes primary microRNAs (pri-miRNAs) which are enzymatically cleaved by the Microprocessor complex, which is composed of Drosha and DiGeorge syndrome critical region 8 (DGCR8) proteins. Processing of the pri-miRNA generates a precursor microRNA (pre-miRNA) that is exported from the nucleus to the cytoplasm. In the cytoplasm, the pre-miRNA is further processed by Dicer, which functions in complex with transactivation response element RNA-binding protein (TRBP) (21–23). Processing of the pre-miRNA by Dicer/TRBP results in the production of a 21-22 nt mature miRNA duplex (24,25). Mature miRNAs function in concert with Argonaute (Ago) proteins to form the RNA-induced silencing complex (RISC) which induces mRNA degradation or translational repression (26,27).

Mature miRNAs are a common ASO target. ASOs that are complementary to the mature miRNA effectively sequester that miRNA within the RISC into a duplex. This sequestration prevents RISC binding to the targeted mRNA, which increases the mRNA lifetime and protein expression. An alternative approach for ASO-based therapeutics involves targeting pre-miRNAs rather than mature miRNAs. ASOs that bind to the apical loop region and block the dicing sites of pre-miR-16, pre-miR-15a, and pre-miR-125b, are effective inhibitors of Dicer/TRBP processing (28). However, ASOs that do not span the dicing site, binding only to the apical loop region, appear to have no effect of the Dicer/TRBP processing of pre-miR-21 (29). We therefore sought to examine the common features of pre-miRNAs that make them suitable targets for ASO steric blocking and cleavage inhibition.

Previously, we found that the junction region of pre-miR-31 is essential for efficient Dicer/TRBP processing and that disrupting junction base pairing inhibits Dicer processing (30). Here, we examined the effect of ASOs that can disrupt the pre-miR-31 junction, promoting an open structure. We found that these junction-disrupting ASOs strongly inhibited pre-miR-31 processing by Dicer/TRBP. Furthermore, we demonstrated that pre-miR-144, which is predicted to contain a similar junction region, is also targetable by ASOs for Dicer/TRBP cleavage inhibition. Examination of the predicted pre-miRNA secondary structures from miRbase revealed that junction regions are a common structural feature among human pre-miRNAs (∼18%). Of the identified junction-containing pre-miRNAs, we tested our steric blocking ASO approach on pre-miR-19a and pre-miR-143 and found these ASOs were effective inhibitors of Dicer/TRBP processing. These results suggest that steric blocking ASOs can be broadly used to inhibit Dicer processing of junction-containing pre-miRNAs. Finally, we demonstrate the effectiveness of these steric-blocking ASOs *in cell* using a dual luciferase reporter assay. These findings provide new insights into our understanding of the role of junction regions in regulating pre-miRNA processing and identifies a new approach for modulating pre-miRNA maturation and translational control.

## MATERIAL AND METHODS

### Preparation of recombinant human Dicer

Human Dicer protein was purified as previously described (31,32) with modifications. Sf9 cells with infected His-tagged Dicer baculovirus was purchased from the University of Michigan protein core. The cell pellet was lysed in ice-cold lysis buffer (50 mM Na_2_HPO_4_ pH = 8.0, 300 mM NaCl, 0.5% Triton X-100, 5% glycerol, 0.5 mM tris(2-carboxyethyl) phosphine (TCEP) and 10 mM imidazole) by sonication. The lysate was pelleted by centrifugation at 30,000 x g for 30 min and the supernatant was mixed with 5 mL Ni-NTA resin (Qiagen) pre-equilibrated with lysis buffer in a 50 mL falcon tube. After gently rocking for 1 h at 4 °C, the resin was pelleted by centrifugation at 183 x g for 10 min. The resin was washed with 45 mL wash buffer (50 mM Na_2_HPO_4_ pH = 8.0, 300 mM NaCl, 5% glycerol, 0.5 mM TCEP and 20 mM imidazole) 5 times and eluted with elution buffer (50 mM Na_2_HPO_4_ pH = 8.0, 300 mM NaCl, 5% glycerol, 0.5 mM TCEP and 300 mM imidazole). The elution was dialyzed against dialysis buffer (20 mM Tris pH = 7.5, 100 mM NaCl, 1 mM MgCl_2_, 0.1% Triton X-100, 50% glycerol) and the purified protein was stored at -80 ℃. Total protein concentration was determined by Bradford assay (Thermo Fisher Scientific) and the concentration of Dicer was quantified using ImageJ (33).

### Expression and purification of TRBP

The expression and purification of human TRBP was based on previously described procedures (34,35). The pET28a-TRBP was purchased from Addgene (Addgene plasmid # 50351). The pET28a-TRBP plasmid was transformed into *E. coil* Rosetta (DE3) pLysS (Novagen), and cells were grown in LB media until the cell density reached an OD_600_=0.6. Protein expression was induced by addition of 0.2 mM Isopropyl β-d-1-thiogalactopyranoside (IPTG) and cells were incubated at 18 ℃ for 20 h. The cells were harvested by centrifugation, resuspended in buffer A (20 mM Tris–HCl pH 8.0, 500 mM NaCl, 25 mM imidazole, and 5 mM β-mercaptoethanol) and lysed by sonication. The lysate was pelleted by centrifugation at 20,000 x g for 30 min and the supernatant was loaded onto a nickel affinity column (Cytiva) and gradient eluted in buffer B (20 mM Tris–HCl pH 8.0, 500 mM NaCl, 250 mM imidazole buffer, and 5 mM β-mercaptoethanol). The protein elution was collected and 10% polyethyleneimine (PEI) was added dropwise (1.5% vol/vol) to remove nucleic acid contaminants. The suspension was stirred at 4 °C for 30 min and the supernatant was collected by centrifugation (30 min at 20,000 rpm). Two sequential ammonium sulfate cuts were performed (with centrifugation for 30 min at 20,000 rpm in between) at 20% and 80% saturation. The pellet from the 80% ammonium sulfate cut was resuspended in buffer C (20 mM Tris-HCl pH 8.0, 2 mM Dithiothreitol (DTT)) and dialyzed against buffer D (20 mM Tris-HCl pH 8.0, 150 mM NaCl, 2 mM DTT) overnight. The dialyzed sample was concentrated and loaded onto a HiLoad 16/600 Superdex 200 column (Cytiva) equilibrated with buffer D.

### Dicer/TRBP complex formation

The purified Dicer and TRBP proteins were mixed at a 1:3 molar ratio and loaded onto a HiLoad 16/600 Superdex 200 column (Cytiva) equilibrated with buffer D. The fractions enriched with complex were pooled, concentrated, and flash frozen in liquid nitrogen for use in processing assays.

### Preparation of DNA templates for *in vitro* transcription

The DNA template for preparation of the HH-pre-miR-31-HDV was generated as previously described (30). The DNA template for preparation of the pre-miR-144, pre-miR-21, pre-let-7c, GG-pre-miR-19a and pre-miR-143 were generated by overlap-extension (OE) polymerase chain reaction (PCR) using EconoTaq PLUS 2x Master Mix (Lucigen) with primers listed in **Table S1.** The OE PCR template for HH-pre-let-7c was digested with EcoRI and BamHI restriction enzymes and inserted into the pUC-19 plasmid. DNA templates for use in *in vitro* transcription reactions were amplified with EconoTaq PLUS 2x Master Mix (Lucigen) using primers UNIV-pUC19_E105 and HH-pre-let7c-AMP-R (**Table S2**).

### Preparation of RNA

RNAs were prepared by *in vitro* transcription in 1× transcription buffer [40 mM Tris base, 5 mM dithiothreitol, 1 mM spermidine and 0.01% Triton-X (pH = 8.5)] with addition of 3–6 mM ribonucleoside triphosphates (NTPs), 10–20 mM magnesium chloride (MgCl_2_), 30–40 ng/μL DNA template, 0.2 unit/mL yeast inorganic pyrophosphatase (New England Biolabs) (36), ∼15 μM T7 RNA polymerase and 10–20% (v/v) dimethyl sulfoxide (DMSO). Reaction mixtures were incubated at 37 °C for 3–4 h, with shaking at 70 rpm, and then quenched using a solution of 7 M urea and 500 mM ethylenediaminetetraacetic acid (EDTA), pH = 8.5. Transcription reactions were boiled for 3 min and then snap cooled in ice water for 3 min. The transcription mixture was loaded onto preparative-scale 10% denaturing polyacrylamide gels for purification. Target RNAs were visualized by UV shadowing and gel bands containing RNA were excised. Gel slices were placed into an elutrap electroelution device (The Gel Company) in 1× TBE buffer. RNA was eluted from the gel at constant voltage (120 V) for ∼24 h. The eluted RNA was spin concentrated, washed with 2 M high-purity sodium chloride, and exchanged into water using Amicon-15 Centrifugal Filter Units (Millipore, Sigma). RNA purity was confirmed on 12% analytical denaturing gels. RNA concentration was quantified via UV-Vis absorbance. Sequences for all RNAs is provided in **Table S3**.

### 32P labeling of RNA

The 5ʹ-end labeling of RNA was performed using 5 pmol of RNA, 1 μL γ-^32^P-ATP (PerkinElmer) and 10 U T4 polynucleotide kinase (New England Biolabs) in a final volume of 10 µL. Before labeling, RNA was boiled for 3 minutes, and snap cooled by placing on ice for another 3 minutes. The radiolabeled RNA was purified on a G-25 column (Cytiva) according to the manufacturer’s instructions. The radiolabeled RNA concentration was determined based on a standard curve which was obtained from the counts per minute of the γ-^32^P-ATP source.

### Dicer/TRBP antisense oligo cleavage assays

Human Dicer/TRBP processing assays were performed as previously described with minimal modifications (30). Concentrated recombinant human Dicer/TRBP protein was diluted in 1× Dicing buffer (24 mM HEPES pH = 7.5, 100 mM NaCl, 5 mM MgCl_2_, 4 μM EDTA). The Dicer/TRBP enzyme was the mixed with 80 U RNaseOUT Recombinant Ribonuclease Inhibitor (Thermo Fisher Scientific), 5× dicing buffer (120 mM HEPES pH = 7.5, 0.5 M NaCl, 25 mM MgCl_2_, 0.02 mM EDTA) and relevant antisense oligo (**Table S4**). The ^32^P-labeled RNA was heated to 95 ℃ for 3 min and then placed on ice for another 3 min. The ^32^P-labeled RNA (1 μL) was added to the Dicer/TRBP mixture (9 μL) and incubated at 37 ℃ for a set period of time. The final RNA and enzyme concentration were 50 nM and 20 nM, respectively. The final antisense oligo concentration was either 500 nM or 5 µM, as indicated in the figures. The reaction was quenched by addition of 10 μL quench buffer (98% Formamide, 20 mM EDTA, trace bromophenol blue and xylene cyanol) at the end time point. The end time points were chosen for each substrate based on a time course study where a substantial fraction of substrate was cleaved (pre-miR-31 at 30 min and pre-miR-144 at 90 min). Experiments were performed in triplicate. The average and standard deviation of the measurements are reported. Significance was determined using one-way ANOVA test.

For the specificity tests, the end time points were 30 min for both pre-miR-21 and pre-let-7c in the pre-miR-31 related assays and 90 min for pre-miR-31 and pre-let-7c in the pre-miR-144 related assays. After the samples were run on 12% denaturing polyacrylamide gels, the gels were exposed to a phosphor screen, which was scanned by a Typhoon Phosphor Imager (GE Healthcare). The gel image was quantified by ImageJ (33). The Dicer/TRBP cleavage ratio was calculated as the sum of the intensity of fully-processed and partially-processed products divided by the sum of the intensity of the fully-processed products, partially-processed products, and remaining substrate (total intensity). Experiments were performed in triplicate. The average and standard deviation of the measurements are reported. Significance was determined using one-way ANOVA test.

### Dicer/TRBP pre-miRNA competition assays

Concentrated recombinant human Dicer/TRBP protein was diluted in 1× Dicing buffer (24 mM HEPES pH = 7.5, 100 mM NaCl, 5 mM MgCl_2_, 4 μM EDTA). The human Dicer/TRBP enzyme complex was pre-mixed with 80 U RNaseOUT Recombinant Ribonuclease Inhibitor (Thermo Fisher Scientific), 5× dicing buffer (120 mM HEPES pH = 7.5, 0.5 M NaCl, 25 mM MgCl_2_, 0.02 mM EDTA) and relevant LNA-ASO (**Table S4**). The RNA mixtures (defined below) were heated to 95 ℃ for 3 min and then placed on ice for another 3 min. One RNA mixture was composed of 500 nM ^32^P-labeled targeted pre-miRNA and 500 nM unlabeled competitive pre-miRNA. In the second RNA mixture, the labeling strategy was inversed. Here 500 nM ^32^P-labeled competitive pre-miRNA was mixed with 500 nM unlabeled targeted pre-miRNA. The RNA mixture (1 μL) was added to a pre-mixed solution of Dicer:TRBP/ASO (9 μL, described above) and incubated at 37℃. The final targeted pre-miRNA, competitive pre-miRNA and enzyme concentration were 50 nM, 50 nM, and 20 nM, respectively. The reactions were conducted in parallel with the two RNA mixtures. The final modified antisense oligo concentration in the reactions was 50, 75, 100, 150, 250, 300, and 400 nM. The reaction was quenched by adding 10 μL quench buffer (98% Formamide, 20 mM EDTA, trace bromophenol blue and xylene cyanol) at the end time point. The end time point for the pre-miR-31 competition assay was 30 minutes and was 90 minutes for the pre-miR-144 competition assay. The samples were run on 12% denaturing polyacrylamide gels. The gels were exposed to a phosphor screen and then scanned by a Typhoon Phosphor Imager (GE Healthcare). The gel image was quantified by ImageJ (33). The Dicer/TRBP cleavage ratio was calculated as the sum of the intensity of fully-processed and partially-processed products divided by the sum of the intensity of the fully-processed products, partially-processed products, and remaining substrate (total intensity). Experiments were performed in triplicate. The average and standard deviation of the measurements are reported. Significance was determined using one-way ANOVA test.

### Dicer/TRBP pre-miRNA SYBR Gold cleavage assays

Human Dicer/TRBP protein processing assays were performed as previously described with minimal modifications (30). Concentrated recombinant human Dicer/TRBP protein was diluted in 1× Dicing buffer (24 mM HEPES pH = 7.5, 100 mM NaCl, 5 mM MgCl_2_, 4 μM EDTA). Dicer/TRBP enzyme was pre-mixed with 80 U RNaseOUT Recombinant Ribonuclease Inhibitor (Thermo Fisher Scientific), 5× dicing buffer (120 mM, pH = 7.5, 0.5 M NaCl, 25 mM MgCl_2_, 0.02 mM EDTA) and antisense oligos. The RNA was heated to 95 ℃ for 3 min and then placed on ice for another 3 min. The RNA (1 μL) was added to the pre-mixed solution of Dicer:TRBP/ASO (9 μL) and incubated at 37 ℃. The final RNA concentrations for pre-miR-144 and pre-miR-143 were 50 nM and 100 nM for GG-pre-miR-19a. The final enzyme concentration was 20 nM for all reactions. The final antisense oligo concentration was either 500 nM or 5 µM for pre-miR-144 and pre-miR-143 and was either 1 µM or 10 µM for GG-pre-miR-19a. The reaction was quenched by adding 10 μL quench buffer (98% Formamide, 20 mM EDTA, trace bromophenol blue and xylene cyanol) at the end time point. The end time point for pre-miR-144 and pre-miR-143 RNA was 120 min and was 30 min for GG-pre-miR-19a based on kinetic analysis of the substrate processing. After the sample was run on a 20% denaturing polyacrylamide gel, the gel was stained with 1× SYBR Gold Nucleic Acid Gel Stain for 30 min (Thermo Fisher Scientific). The gel image was quantified by ImageJ (33). The Dicer/TRBP cleavage ratio was calculated as the sum of the intensity of fully-processed and partially-processed products divided by the sum of the intensity of the fully-processed products, partially-processed products, and remaining substrate (total intensity). The individual fully-processed 5p strand and 3p strand intensity ratios were calculated as the sum of the intensity of fully-processed 5p or 3p strand intensity divided by the sum of the intensity of the fully-processed products, partially-processed products, and remaining substrate. Experiments were performed in triplicate. The average and standard deviation of the measurements are reported. Significance was determined using one-way ANOVA test.

### Cataloging of junction regions of human pre-miRNAs

All 1,613 pre-miRNA secondary structures were analyzed based on the structure provided in the miRbase human microRNA database (37). We categorized pre-miRNAs as junction-containing or not based on the following criteria (**Fig. S1**). First, we evaluated if there were predicted base pairs between the dicer cleavage site and the apical loop. If one or both cleavage sites were within that base paired region, those pre-miRNAs were classified as not containing a junction region. All other pre-miRNAs would be considered as containing a junction if one of the two additional criteria were met; a) at least one cleavage site is present in a single stranded region with at least one unpaired nt below the cleavage site, and b) if both cleavage sites are within a base paired region, an unpaired region exists between two cleavage sites. All relevant information, including the cleavage site position (junction region, stem region, apical loop or internal loop), and the size and composition of the base paired junction region, was recorded and analyzed.

### pmirGLO plasmid construction

The pmirGLO plasmid construction was based on the manufacturer manuals. The pmirGLO dual-luciferase miRNA target expression vector was purchased from Promega. The vector was digested with PmeI and XbaI restriction enzymes (New England Biolabs). The inserts for pmirGLO miR144 and pmiRGLO miR31 plasmids were purchased from Integrated DNA Technologies (**Table S5**) and were also digested with PmeI and XbaI restriction enzyme. The inserts were each ligated with digested pmirGLO plasmids using a quick ligasation kit (New England Biolabs) and transformed into JM109 competent cells (Promega). The plasmid was verified by digestion with the NotI restriction enzyme (New England Biolabs), according to manual.

### pCMV plasmid construction

The pCMV miR144 plasmid was a generous gift from Dr. Eric Lai (Memorial Sloan Kettering Cancer Center). The insert for pCMV miR31 was generated by OE PCR using primers listed in **Table S6**. The insert was digested by XhoI and BglII (New England Biolabs) and ligated with digested pCMV vector. The ligated product was transformed into DH5α competent cells, and the plasmid integrity was verified by sequencing.

### *In cell* dual luciferase assays

HEK293 cells (ATCC) were grown in Dulbecco’s Modified Eagle Medium (DMEM, Gibco) with 10% fetal bovine serum (FBS, Gibco) and 1% Penicilin/Streptomycin. HEK293 cells were detached by trypsin treatment and seeded into 96-well plates with 40K cells per well. Cells were allowed to recover for about 24 h. Subsequently, these cells were co-transfected with 50 ng pmirGLO plasmid (reporter plasmid), 50 ng pCMV plasmid (overexpression plasmid), and 10 nM final concentration of ASO or positive/negative control. Positive control ASOs were purchased from Horizon Discovery. Post transfection, the plates were incubated for another 24 h. The Promega dual-luciferase reporter assay system was used to detect firefly and *Renilla* luciferase activities and the data was analyzed according to the manufacturer protocols.

### Quantitative Real-Time PCR Analysis

HEK293 cells (ATCC) were grown in DMEM (Gibco) with 10% FBS (Gibco) and 1% Penicilin/Streptomycin. HEK293 cells were detached by trypsin treatment and seeded into 24-well plates with 200K cells per well. Cells were allowed to recover for about 24 h, and then co-transfected with 250 ng pCMV plasmid (overexpression plasmid) and 10 nM final concentration of antisense oligo or positive/negative control. Post transfection, the plates were incubated for another 24 h. Cells were detached by trypsin treatment and RNAs were extracted using miRNeasy Mini Kit (Qiagen). RNA concentration was determined by NanoDrop One C (ThermoFisher Scientific). Expression level of all microRNAs was assessed utilizing the Taqman advanced miRNA assays (Applied Biosystems) and expression level of U6 level was assessed utilizing the Taqman microRNA assay (Applied Biosystems). The complementary DNA (cDNA) was reverse transcribed by Taqman Reverse Transcription Kit (Applied Biosystems) for U6 and Taqman advanced miRNA cDNA synthesis Kit (Applied Biosystems) for miRNAs. U6 was used as internal control for all miRNA levels. The Ct was determined using single threshold method. All experiments were performed in triplicate, with all samples normalized to U6 RNA level and relative expression levels were calculated using 2^-ΔΔCt^ method.

## RESULTS

### Antisense oligos inhibit pre-miR-31 cleavage by Dicer/TRBP

RNAs are known to be conformationally dynamic molecules, and the conformational dynamics of pre-miRNAs can be important for regulating Dicer/TRBP binding and processing. Studies on pre-miR-21 revealed the existence of both ground and excited state conformations and demonstrated that Dicer cleaved the excited state conformation more efficiently than the ground state conformation (38). Pre-miR-21 processing can also be inhibited by drug-like small molecules, which bind to pre-miR-21 and stabilize it into a conformation that is a poor substrate for Dicer processing (35). We previously identified the junction region is a regulatory element in pre-miR-31 and demonstrated that altering the stability of this region affected Dicer/TRBP cleavage (**Fig. 1a**) (30). The open structure of pre-miR-31, which does not contain a pre-formed junction region, is a poor substrate for Dicer/TRBP cleavage (30). We therefore explored the use of steric-blocking antisense oligos that could compete with pre-miR-31 junction and stabilize the RNA into an “open” conformation to inhibit pre-miR-31 maturation.

**Figure 1.**
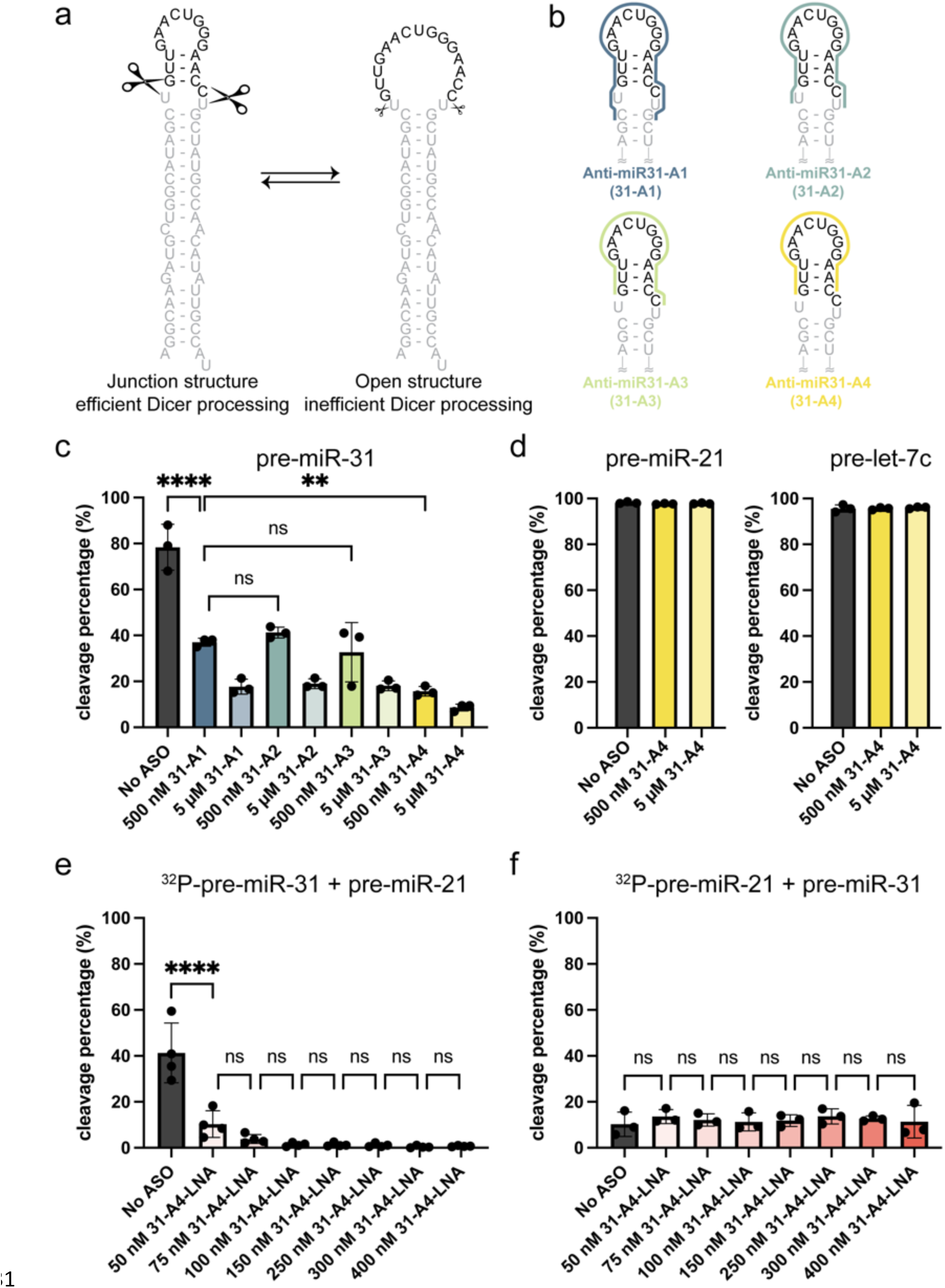
Antisense oligos that target the pre-miR-31 junction structure are potent and specific inhibitors of Dicer/TRBP processing *in vitro*. **a)** Pre-miR-31 conformational equilibrium highlighting the junction structure containing structure, a good substrate for processing by the Dicer/TRBP complex, and an open loop structure, which is a poor substrate for Dicer/TRBP processing. The mature miR-31 duplex sequence is gray and the cleaved off apical loop is black. Scissors indicate Dicer cleavage sites with sizes to reflect the extent to which each conformer is processed. **b)** Pre-miR-31 ASO design. Colored lines span the nucleotides within pre-miR-31 that each anti-miR-31 ASO is complimentary to. **c)** Effect of addition of anti-miR-31-A1-A4 to a pre-miR-31 Dicer/TRBP cleavage assay. **d)** Dicer/TRBP processing assays of pre-miR-21 and pre-let-7c in the presence and absence of anti-miR31-A4. **e)** Competition cleavage assay conducted on ^32^P-pre-miR-31 and unlabeled pre-miR-21 in the presence of anti-miR-31-A4-LNA. **f)** Competition cleavage assay conducted on ^32^P-pre-miR-21 and unlabeled pre-miR-31 in the presence of anti-miR-31-A4-LNA. For all processing assays, the average and standard deviation from n=3 or n=4 independent assays are presented. **** p<0.0001, **p<0.01, and ns indicates no significant difference from an ordinary one-way ANOVA Tukey analysis.

We designed a series of antisense oligos, intended to bind to the pre-miR-31 apical loop region and disrupt the junction base pairs (**Fig. 1b, Table S4**). Anti-miR31-A1 (31-A1) is the longest ASO, spanning the apical loop, junction, dicing site, and disrupting one additional C-G base pair directly beneath the internal loop. Anti-miR31-A2 (31-A2) is complementary to full apical loop sequence of the open conformation for pre-miR-31, which partially sequesters the Dicer cleavage site in the loop. Anti-miR31-A3 (31-A3) binds to the apical loop sequence of the pre-miR-31 without spanning the Dicer cleavage site (i.e. the sequence that would be cleaved by Dicer). Anti-miR31-A4 (31-A4) is the shortest ASO and only disrupts the junction base pairs, annealing across the apical loop. We expected all ASOs to disrupt junction base pairing, promoting intermolecular rather than intramolecular interactions at the junction region. *In vitro* Dicer/TRBP processing assays on pre-miR-31 in the absence or presence of different concentrations of various Anti-miR31 ASOs revealed significant inhibition of processing with 31-A4 providing the highest degree of inhibition (**Fig. 1c**). Subsequent studies were therefore conducted only with 31-A4.

We next used the siRNA Target Finder tool (https://www.genscript.com/tools/sirna-target-finder) to determine if 31-A4 would target any other pre-miRNA sequence. No pre-miRNAs were identified that had sequences complementary to 31-A4. To examine the specificity of 31-A4 inhibition, we assessed the processing of pre-miR-21 and pre-let-7c incubated with excess amount of 31-A4. We did not observe a detectable reduction in Dicer/TRBP processing for either pre-miR-21 or pre-let-7c in the presence of 31-A4 ASO (Fig. 1d). These results suggest that 31-A4 acts specifically to inhibit pre-miR-31 processing and is not a global repressor of Dicing for pre-miRNAs.

While our *in vitro* assays were conducted with DNA oligonucleotides containing a native phosphodiester backbone, in the cell, naked DNAs are readily degraded and are therefore not suitable for antisense oligo design (1). A variety of sugar, backbone, and base modifications can be incorporated into the ASO to enhance the ASO stability and binding affinity (1,16,39,40). We therefore ordered a new version of the 31-A4 ASO that contained a modified phosphorothioate backbone and mixed locked nucleic acid (LNA) bases where the 2’ oxygen and the 4’ carbon of some pentose sugars were linked with a methylene bridge (anti-miR31-A4-LNA, 31-A4-LNA, **Table S4**). We employed a competition assay to assess the impact of our modified ASO on the Dicer/TRBP processing of pre-miR-31 in the presence of a competitor pre-miRNA. Here, equal amounts of the target pre-miRNA (pre-miR-31) and a competitor pre-miRNA (pre-miR-21) were mixed; one of the pre-miRNAs was radiolabeled at the 5’-end while the other pre-miRNA was unlabeled. Incubation of the pre-miRNA mixture in the presence and absence of 31-A4-LNA and addition of Dicer/TRBP permits the analysis of the radiolabeled pre-miRNA. By changing which pre-miRNA was radiolabeled, we were able to detect the cleavage percentage of each pre-miRNA in the competition cleavage assay. We found that pre-miR-31 cleavage was inhibited by 31-A4-LNA in a dose dependent manner (Fig. 1e) while there was no detectable impact on pre-miR-21 cleavage by Dicer/TRBP (Fig. 1f). Collectively, we found that all junction-spanning ASOs of pre-miR-31 strongly inhibit Dicer/TRBP cleavage with no off-target effects on other pre-miRNAs examined.

### Pre-miR-144 is a targetable pre-miRNA containing a junction region

Secondary structure probing of pre-miR-144 revealed a junction-containing structure (41). Mutants that disrupted the pre-miR-144 junction base pairs inhibited Dicer processing *in cell* (42), similar to our observations in pre-miR-31 (30). We therefore designed steric blocking ASOs to target the pre-miR-144 junction region and assessed the effect of the ASOs on inhibiting its Dicer-mediated maturation.

Using the secondary structure of pre-miR-144 as a reference (Fig. 2a), we designed four ASOs based on similar principles described above for pre-miR-31 (Fig. 2b). Anti-miR144-A1 (144-A1) was designed to fully destabilize the junction, span the dicing site, and destabilize one additional A-U base pair positioned directly beneath the 1×2 internal loop. Anti-miR144-A2 (144-A2) spans the internal loop, junction region, and apical loop while anti-miR144-A3 (144-A3), and anti-miR144-A4 (144-A4) are shorter ASOs that destabilize the junction base pairs, locking the junction region in an “open” conformation. In our *in vitro* processing assays, all pre-miR-144 ASOs inhibited Dicer/TRBP cleavage to a similar extent (Fig. 2c). The 144 ASOs did not impact the processing of other pre-miRNAs by Dicer/TRBP (**Fig. S2a, b**), even for the pre-miR-31, which also contains a junction region.

**Figure 2.**
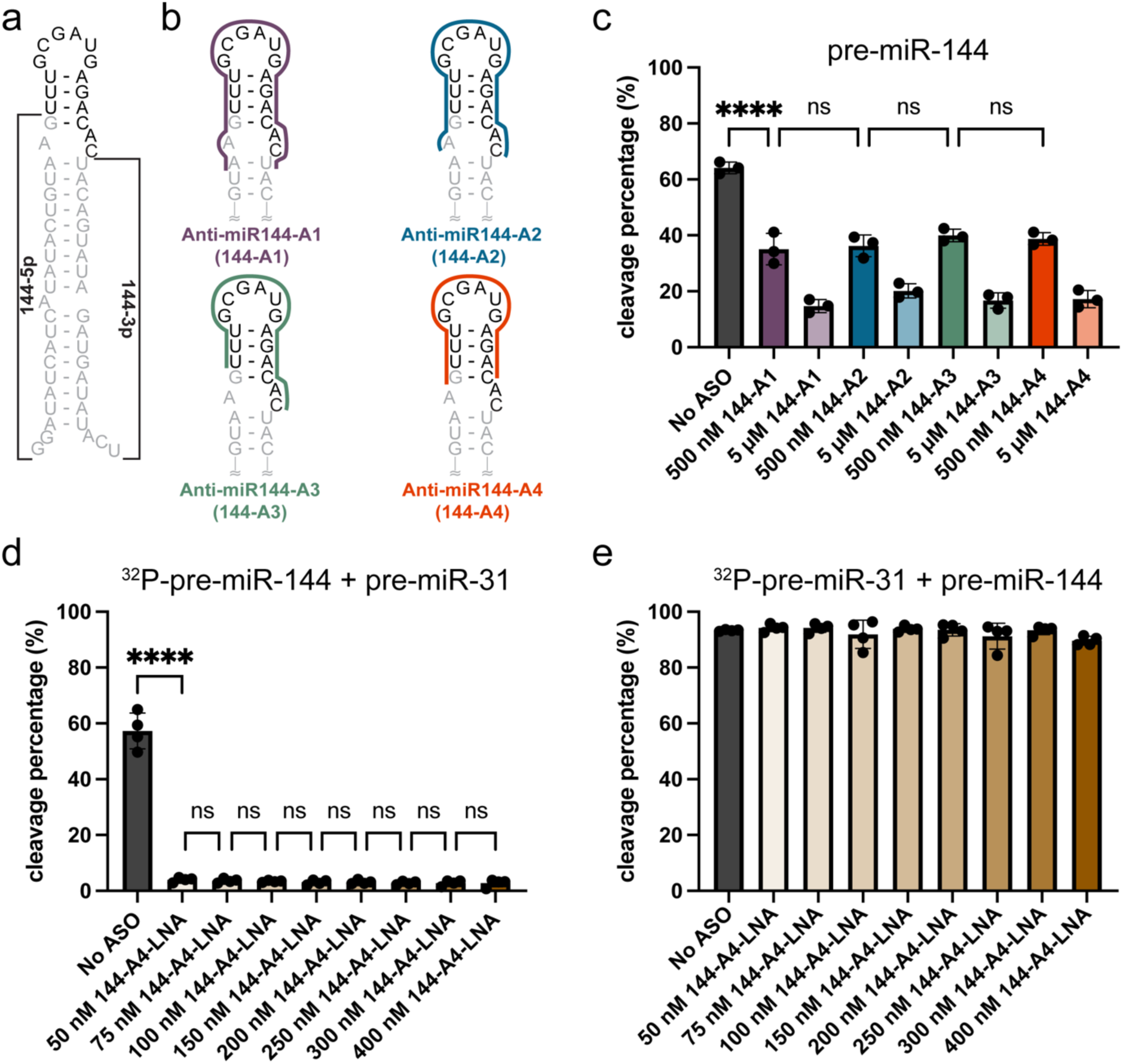
ASOs targeting the pre-miR-144 junction inhibit Dicer/TRBP processing *in vitro*. **a)** Predicted secondary structure of pre-miR-144. The mature miR-144 duplex sequence is gray and the cleaved apical loop is black. The 144-5p and 144-3p products are labeled, 144-3p serves as the guide strand. **b)** ASO design. Colored lines span the nucleotides within pre-miR-144 that each anti-miR-144 ASO is complimentary to. **c)** Effect of addition of anti-miR-144-A1-A4 to a pre-miR-144 Dicer/TRBP cleavage assay. **d)** Competition cleavage assay conducted on ^32^P-pre-miR-144 and unlabeled pre-miR-31 in the presence of anti-miR-144-A4-LNA. **e)** Competition cleavage assay conducted on ^32^P-pre-miR-31 and unlabeled pre-miR-144 in the presence of anti-miR-144-A4-LNA. For all processing assays, the average and standard deviation from n=3 or n=4 independent assays are presented. **** p<0.0001 and ns indicates no significant difference from an ordinary one-way ANOVA Tukey analysis.

Because all the anti-pre-miR-144 ASOs that we tested inhibited Dicer processing to a similar extent, we moved forward with the 144-A4 construct, the minimal ASO construct that only disrupts the junction formation. The guide strand is the functional component of the mature miRNA duplex which forms the RISC and promotes sequence-specific binding to the target mRNA. The production of the guide strand therefore plays an important role in post-transcriptional gene expression. In pre-miR-144, the guide strand is the 3ʹ-end cleavage product (3p, Fig. 2a), which is masked in our 5ʹ-^32^P radiolabeled experiments. To ensure that the ASO didn’t differentially impact generation of the 5ʹ (5p) or 3ʹ (3p) product, we performed Dicer/TRBP cleavage assays and analyzed all processing products using SYBR gold stain (**Fig. S3**). Consistent with our observations from the 5ʹ-^32^P radiolabeled processing experiments, we observed that addition of 144-A4 inhibited the production of mature miR-144 duplex when analyzed by SYBR staining (Fig. 2a). Furthermore, we analyzed the production of the miR-144 3p and 5p strands individually. Here we found a similar dose-dependent inhibition of 5p and 3p products in the presence of 144-A4, indicating that the production of both strands of mature miR-144 was inhibited to a similar extent (**Fig. S3b, c**). Unlike using a radiolabeled substrate, the band intensity of SYBR stained gels is dependent on RNA size, therefore we cannot make direct comparisons across samples. Importantly, these findings ensured that both strands of pre-miR-144 were produced and the processing of the 3p strand was also inhibited by addition of 144-A4.

We next examined the effect of a modified anti-miR144-A4 ASO (144-A4-LNA) on Dicer/TRBP processing. Competition cleavage assays were conducted on pre-miR-144 in the presence of pre-miR-31, which contains a similar junction region. We observed strong cleavage inhibition of pre-miR-144 by 144-LNA (Fig. 2d) while pre-miR-31 cleavage levels were unaffected (Fig. 2e), indicating that the pre-miR-144 ASO can strongly inhibit pre-miR-144 maturation without perturbing the processing of other similarly structured pre-miRNA.

### The junction region is a common structural feature within pre-miRNAs

We previously showed that the stability of the junction region is important for pre-miR-31 processing (30). Furthermore, we demonstrated that the junction region is a targetable element for inhibiting Dicer/TRBP cleavage of pre-miR-31 (Fig. 1) and pre-miR-144 (Fig. 2) using ASOs. We were therefore interested to know how many pre-miRNAs are predicted to contain a junction region. We examined the predicted secondary structures of pre-miRNAs provided in the miRbase database (37,43–46) and classified pre-miRNAs based on whether or not they are predicted to contain a junction (**Fig. S1**). Among the 1,663 pre-miRNAs, 296 (∼18%) are predicted to contain a junction region (Fig. 3a). Of these 296 pre-miRNAs, ∼70% are predicted to contain junction regions between three and five base pairs in length (Fig. 3b).

**Figure 3.**
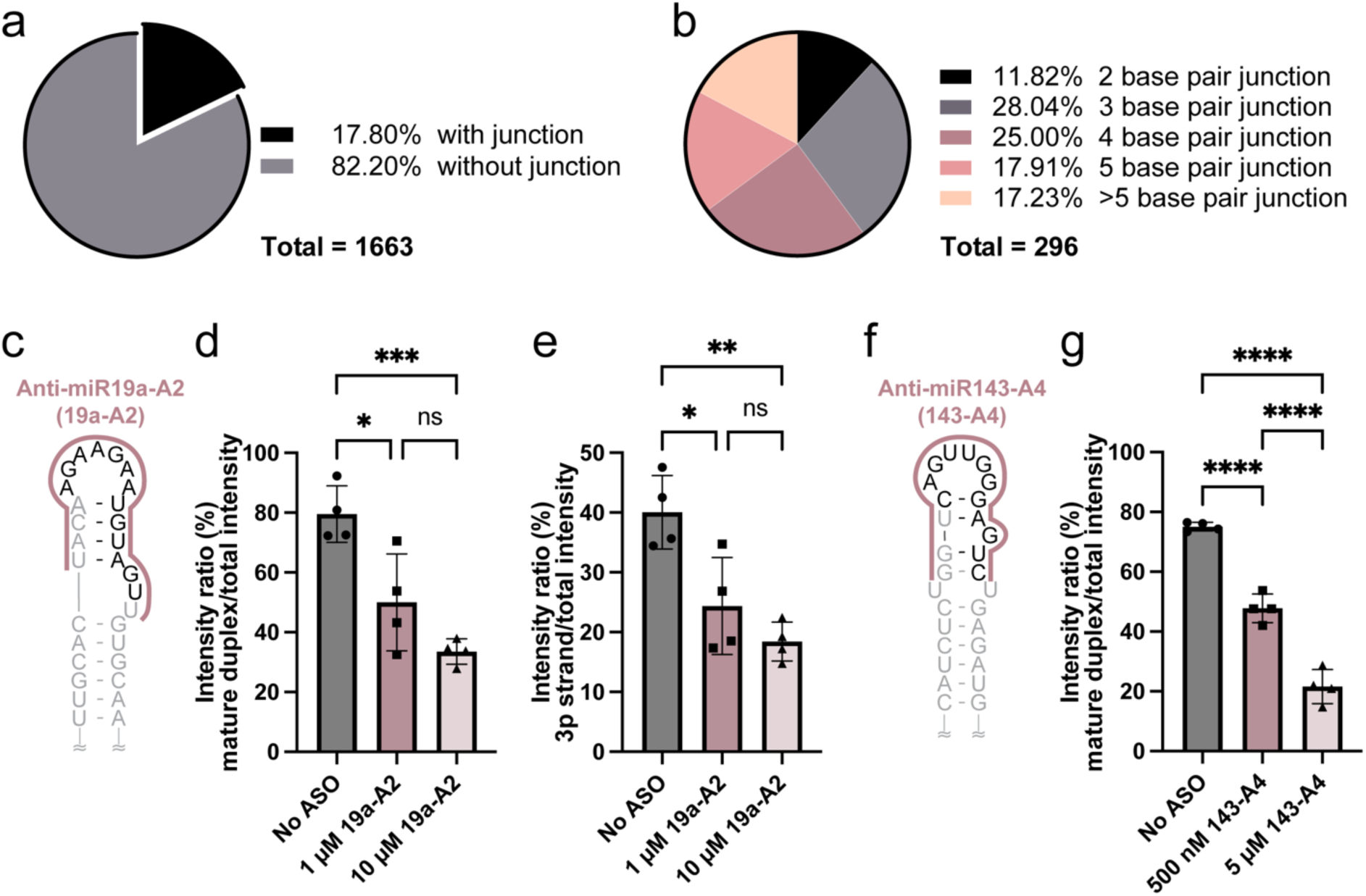
Junction regions are a common feature of pre-miRNAs and are effective ASO targets. **a)** Analysis of secondary structures predicted in the miRbase reveals the prevalence of junction-containing pre-miRNAs. **b)** Breakdown of predicted junction length. **c)** Pre-miR-19a is predicted to contain a four base pair junction. Anti-miR19a-A2 binding site is indicated. The mature miR-19a duplex sequence is gray and the cleaved apical loop is black. **d)** Quantification of pre-miR-19a processing products with increasing concentrations of anti-miR19a-A2. **e)** Quantification of the pre-miR-19a 3p strand product in Dicer/TRBP processing assays that include increasing concentrations of anti-miR19a-A2. **f)** Pre-miR-143 predicted secondary structure with the anti-miR143-A4 binding site indicated. The mature miR-143 duplex sequence is gray and the cleaved off apical loop is black. **g)** Quantification of pre-miR-143 processing products with increasing concentrations of anti-miR143-A4. For all processing assays, average and standard deviation from n=4 independent assays are presented. **** p<0.0001, ***p<0.001, **p<0.01, *p<0.05, and ns indicates no significant difference from an ordinary one-way ANOVA Tukey analysis.

### Targeting junction containing pre-miRNAs is a reasonable approach for inhibiting Dicer/TRBP processing

To expand our study of ASOs targeting junction-containing pre-miRNAs, we conducted additional processing assays using pre-miR-19a and pre-miR-143, two pre-miRNAs predicted to contain four base pair junction based on our analysis. We designed two antisense oligos, anti-miR19a-A2 (19a-A2) and anti-miR143-A4 (143-A4), which were designed to target pre-miR-19a and pre-miR-143, respectively (Fig 3c, f). To simplify the transcription and purification of pre-miR-19a, the first adenosine was replaced with a guanosine to generate GG-pre-miR-19a, which has the same predicted secondary structure as the native pre-miR-19a (**Fig. S4**). Due to the high A U content within the junction and apical loop of pre-miR-19a, we designed a longer A4-like ASO, spanning the dicer cleavage site, junction region, and internal and apical loops, to ensure tight binding under the reaction conditions. Incubation with 10-fold and 100-fold excess ASO resulted in significant inhibition of Dicer/TRBP processing of both GG-pre-miR19a, and pre-miR-143 (Fig. 3c**-g**). The guide strand (3p) intensity ratio for GG-pre-miR-19a was also significantly reduced when treated with high concentrations of 19a-A2 (Fig 4e), as the intact GG-pre-miR-19a accumulates (**Fig. S5**). For pre-miR-143, we detected similar inhibitory effects when treated with a high concentration of 143-A4 (Fig. 3g**, S6**).

**Figure 4.**
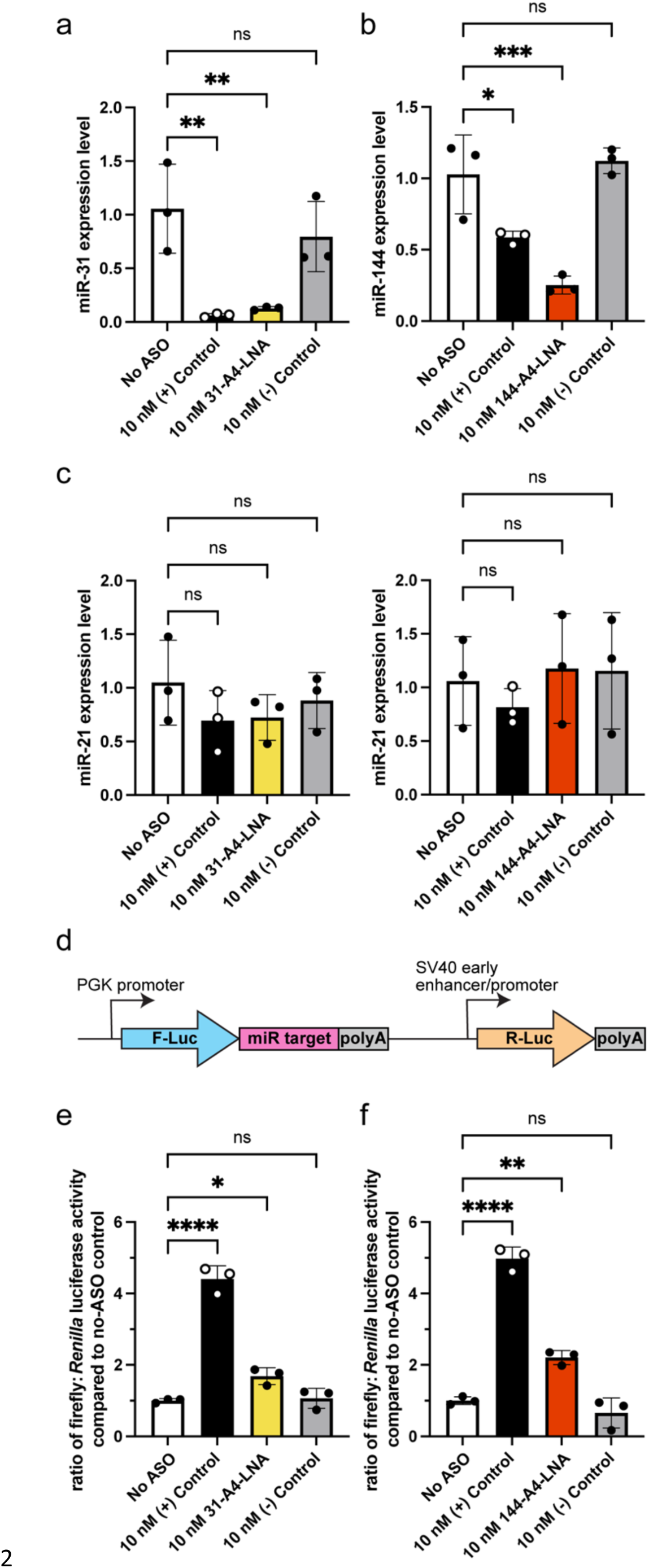
Anti-junction-region ASOs can inhibit pre-miRNA maturation *in cell*. **a)** RT-qPCR analysis of mature miR-31 expression levels in HEK293 cells upon treatment with various ASOs. **b)** RT-qPCR analysis of mature miR-144 expression levels in HEK293 cells upon treatment with various ASOs. **c)** RT-qPCR analysis of mature miR-21 expression levels in HEK293 cells within miR-31 (left) and miR-144 (right) *in cell* experiment samples. **d)** Diagram of the pmirGLO vector for the dual luciferase assays. The expression of firefly luciferase (F-Luc, blue) is under the control of the target mature miRNA while *Renilla* luciferase (R-Luc, yellow) expression is not controlled by any miRNA sequence. R-Luc luciferase signals are used as internal normalization control. **e,f)** Luciferase reporter assay for miR-31 (**e**) and miR-144 (**f**) with various ASO treatment in HEK293 cells. The post-transcriptional inhibition of F-Luc translation by miR-31 and miR-144 are recovered by anti-junction-region ASOs, suggesting that these ASO function to reduce the levels of miR-31 and miR-144. For all assays, average and standard deviation from n=3 independent assays are presented. **** p<0.0001, **p<0.01, *p<0.05, and ns indicates no significant difference from an ordinary one-way ANOVA analysis.

While the molecular details of the interaction between the human Dicer helicase domain and the apical loop of pre-miRNAs is masked in human Dicer-RNA complex structures, this domain provides important contacts that mediate the specific processing of pre-miRNAs (47–49). We therefore sought to determine if the inhibition of Dicer processing observed in our ASO assays is the result of disruption between the pre-miRNA apical loop and Dicer helicase domain recognition. We selected pre-let-7c from the pool of non-junction containing pre-miRNAs to investigate this possibility. We designed anti-let-7c-A1 (**Fig. S7a**, **Table S4**), a 14-nt ASO designed to target the apical loop region of the non-junction-containing pre-let-7c. Interestingly, we found that even 100-fold excess of anti-let-7c-A1 did not significantly inhibit Dicer/TRBP cleavage on pre-let-7c (**Fig. S7b**), suggesting that the ASO does not function as a steric block to Dicer/TRBP recognition. Collectively, our findings suggest that, for junction-containing pre-miRNAs, targeting of the junction and apical loop regions is an alternative approach for inhibition of Dicer processing. This success suggests that the junction region is an important domain within a subset of pre-miRNAs, and that junction formation facilitates the efficient cleavage by Dicer/TRBP.

### ASO targeting of junction-containing pre-miRNAs inhibit Dicer cleavage *in cell*

Our *in vitro* processing assays revealed that ASOs targeting the junction region of pre-miRNAs were inhibitors of Dicer/TRBP processing. We next sought to validate the inhibitory effects of 31-A4-LNA and 144-A4-LNA on Dicer/TRBP processing of their respective pre-miRNA elements *in cell*. First, we quantified mature miRNA levels (miR-31 or miR-144) in HEK293 cells after transfection with the relevant ASO using quantitative reverse transcription polymerase chain reaction (qRT-PCR). We detected significant inhibition of mature miR-31 levels in cells transfected with either commercially available positive controls (complementary to the mature miRNA sequence) or our 31-A4-LNA (Fig. 4a). We did not detect significant changes in mature miR-31 levels in cells transfected with a negative control (**Table S4**). Transfection of HEK293 cells expressing pre-miR-144 with 144-A4-LNA also strongly inhibited miR-144 production (**Fig.4b**). Importantly, we verified that ASO transfection did not affect off-target miR-21 levels (Fig. 4c).

Both 31-A4-LNA and 144-A4-LNA ASOs were effective in reducing mature miRNA levels. We therefore used a dual luciferase assay to determine if the ASOs affected gene expression (Fig. 4d). We co-transfected an overexpression plasmid (for either pre-miR-31 or pre-miR-144), relevant ASOs, and the respective dual luciferase reporter plasmid into HEK293 cells. After incubation for 24 hours, we quantified the relative luciferase activity. We found that the luciferase signal was elevated in both 31-A4-LNA and 144-A4-LNA treated samples, relative to their No ASO treated controls (Fig. 4e**,f**). These results further confirm that our junction region targeted ASOs inhibit miRNA production *in cells*.

## DISCUSSION

ASOs play important roles in nucleic therapeutics and there is growing interest in developing novel ASOs that target different aspects of nucleic acid biology (1,50). MicroRNAs are one type of nucleic acid that have been targeted by ASOs for therapeutic purposes (51,52). Miravirsen is an ASO that inhibits hepatitis C virus (HCV) replication in liver cells by disrupting the interaction between miR-122 and sequences in the 5ʹ-UTR of the viral genome (53,54). MRG-110 targets miR-92a, a negative regulatory miRNA in cardiovascular diseases, to improve angiogenesis (55). A number of miRNA-targeted ASOs, which have demonstrated effectiveness *in vitro* are currently in preclinical trial or clinical trials for further efficacy and safety assessment (56). However, most ASOs target the mature sequence of microRNAs (56), with relatively little research aimed at targeting pri/pre-element microRNA using ASOs (28). Interestingly, a virus-encoded miRNA (HHV-6A miR-aU14) has been shown to inhibit pre-miR-30 processing through RNA-RNA interactions within the pre-miR-30 apical loop (57). However, the artificial ASOs, specifically designed to inhibit pre-miRNA processing, have generated mixed results (28,29).

Many families of RNAs require active structures for biological activity, and misfolding of the structure can trap an RNA in an inactive functional state. The ability of ASOs to invade RNA structures and induce their misfolding has been examined in several systems. The group I self-splicing intron from *Candida albicans* is positioned within the ribosomal RNA (rRNA) precursor (58,59) and the self-splicing of this sequence is essential for ribosome maturation (60). Because the structure of rRNAs is essential for ribosome function (61), inducing misfolding of the group I intron therefore affects the processing of the rRNA precursor which can ultimately inhibit ribosome assembly. In this manner, ASOs designed to invade the functional structure of group I intron and trap the P3 stem of the intron into a misfolded state are effective inhibitors of self-splicing in *C. albicans* (58).

ASO-based structure invasion has also been demonstrated to impact microRNA biogenesis. As mentioned above, miravirsen sequesters mature miR-122 and inhibits HCV replication (54). However, the sequestration of miR-122 may not be the only factor precipitating the observed inhibition of replication. Studies also showed that the pre-miR-122 structure was invaded by miravirsen and that the addition of miravirsen inhibited the processing of pre-miR-122 *in cell* (62). In a similar manner, miravirsen was also able to invade the pri-miR-122 stem-loop structure, inhibiting Drosha processing of the pri-element, which added a second layer of inhibition to the miRNA biogenesis pathway (62). These studies lay the foundation for the development of ASOs that invade the structures of functional RNAs as therapeutic method.

Our previous research uncovered the importance of the pre-miR-31 junction region in regulating Dicer/TRBP processing (30). We found that the junction-containing structure was a good substrate for Dicer/TRBP processing while mutations that disrupted the junction base pairs reduced the processing by Dicer/TRBP. Therefore, pre-miR-31 can be used as a model for investigating if ASOs that invade and disrupt the junction structure are potent inhibitors of Dicer/TRBP processing.

We found that addition of ASOs that blocked junction base pair formation within pre-miR-31 and locked the RNA into an open junction conformation strongly inhibited Dicer/TRBP processing. These junction-targeting ASOs are specific for inhibiting pre-miR-31 processing and do not broadly inhibit Dicer/TRBP cleavage of other pre-miRNAs. ASOs that were composed of chemically-modified bases and backbone linkages strongly inhibited pre-miR-31 cleavage in the presence of other pre-miRNAs both *in vitro* and *in cell*.

The utility of junction-targeting ASOs was not limited to pre-miR-31. Pre-miR-144 has a similar junction structure and we demonstrated that ASOs designed to disrupt that junction structure were both highly specific and efficient in inhibiting Dicer/TRBP cleavage. Our bioinformatics analysis revealed that ∼20% of human pre-miRNAs are predicted to contain junction regions, and therefore may by targetable using this antisense oligo approach. Previous studies using ASOs that targeted the apical loop of pre-miR-21 had no impact on Dicer/TRBP cleavage (29). Interestingly, unlike pre-miR-31 and pre-miR-144, pre-miR-21 does not contain a junction structure. The presence of a junction region in both pre-miR-144 and pre-miR-31 may explain our success in inhibiting the processing of these pre-miRNAs with ASOs that invade the junction structures. These findings emphasize the importance of biomolecule structure in the design and development of drug-like molecules.

Anti-miRNA ASOs target the guide strand within the RISC complex and interrupt the recognition between RISC and the mRNA 5ʹ-UTR (Fig. 5A). In contrast to the conventional anti-miRNA ASO design strategy, our approach uses ASOs to invade the junction region of pre-miRNAs which alters the structure of the pre-miRNA and ultimately inhibits mature miRNA production by reducing Dicer processing of the substrate (Fig. 5B). These structure-invading ASOs function at an upstream step in the miRNA biogenesis pathway and therefore serve as a complementary approach to controlling gene expression relative to conventional anti-miRNA ASOs. Collectively, these studies add to our understanding about the regulatory role of junction structure in Dicer/TRBP processing.

**Figure 5.**
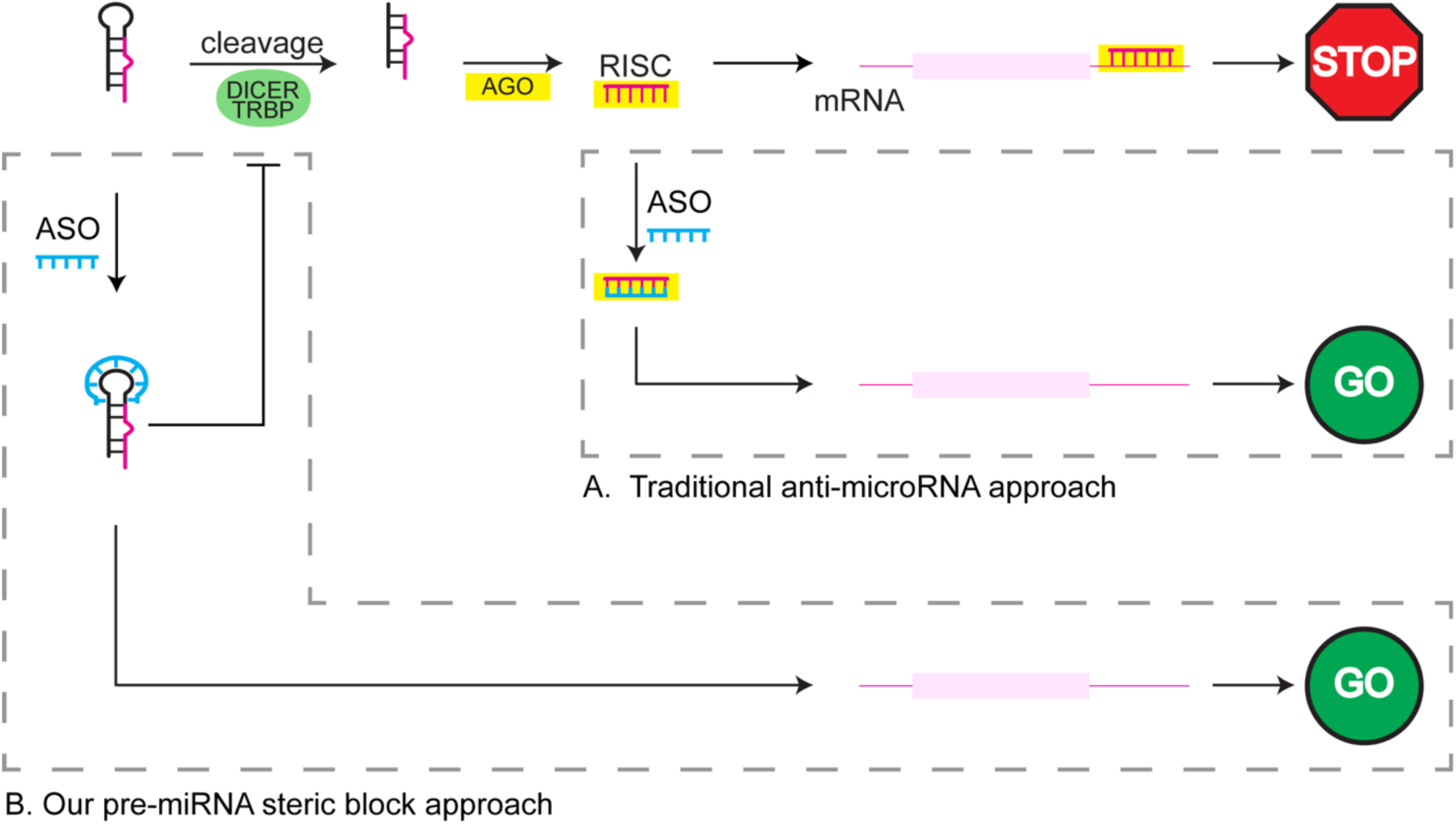
Junction-targeting ASOs function at an upstream step relative to anti-miRNA ASOs. In the cytoplasm, pre-miRNAs are cleaved by the Dicer/TRBP complex to generate a miRNA duplex. The binding of the duplex and selection on the guide strand by AGO forms a functional RNA-induced silencing complex (RISC). RISC binding to the 3’ UTR of target mRNA reduces protein expression. Traditional anti-miRNA ASOs (A, upper right box) interrupt this RISC-mRNA binding to enhance protein expression by sequestering the guide strand inside the RISC in a duplex state. Our strategy (B, lower left box) uses ASOs that target pre-miRNAs containing junction regions to inhibit Dicer/TRBP processing which further reduces the mature miRNA levels in cells. The reduced mature miRNA levels further downregulate function RISC levels to enhance protein expression.

## AUTHOR CONTRIBUTIONS

Sicong Ma: Conceptualization, Investigation, Formal analysis, Project administration, Validation, Writing—original draft, Writing—review & editing. Samantha Howden: Investigation. Sarah C. Keane: Conceptualization, Formal analysis, Funding acquisition, Project administration, Supervision, Writing—review & editing.

## ACKNOWLEDGEMENTS

We are grateful to Dr. Matthew Soellner and members of his lab, specifically Nathalie M Vandecan, for training and guidance in the implementation of the cell-based assays. Additionally, we thank Dr. Eric Lai for providing the pCMV plasmid for pre-miR-144 expression. We thank members of the Keane Lab for helpful discussion.

