## Supplementary Data for "Use of steric blocking antisense oligonucleotides for the targeted inhibition of junction containing precursor microRNAs"

**Table S1. Overlap extension PCR primers for generation of pre-miRNA templates.**

| Construct | Primer name | 5'-sequence-3'[a] |
|---|---|---|
| HH-pre-miR-21 | HH-pre-miR-21-1F | TAATACGACTCACTATAGGCTCG |
| | HH-pre-miR-21-2R | ACCCTGATGGTGTCTGAAAAACGTACCCTGATGGTGTACGAGCCTATAGTGAGTCGTAT |
| | HH-pre-miR-21-3F | TTCAGACACCATCAGGGTCTGCTGATAAGCTACTGATGAGTCCGTGAGG |
| | HH-pre-miR-21-4R | CTAGACGGTACCGGGTACCGTTTCGTCCTCACGGACTCATCAGTA |
| | HH-pre-miR-21-5F | ACCCGGTACCGTCTAGCTTATCAGACTGATGTTGACTGTTGAATCTCATGGCAACACCAG |
| | HH-pre-miR-21-6R | mGmACAGCCCATCGACTGGTGTTGCCATGAGATT |
| pre-miR-let7c | pre-miR-let7c-nat-1F | CCGGAATTCTAATACGACTCACTATAGGGCTCGTACACCATCAGGGTACGTTTTTCA |
| | pre-miR-let7c-nat-2R | GCAGACCCTGATGGTGTCTGAAAAACGTACCCTGATGGTGTACGAGCC |
| | pre-miR-let7c-nat-3F | ACACCATCAGGGTCTGCTACTACCTCACTGATGAGTCCGTGAGGACGAA |
| | pre-miR-let7c-nat-4R | CCTCAGACGGTACCGGGTACCGTTTCGTCCTCACGGACTCAT |
| | pre-miR-let7c-nat-5F | CCGGTACCGTCTGAGGTAGTAGGTTGTATGGTTTAGAGTTACACCCTGGGAGTTAACTGT |
| | pre-miR-let7c-nat-6R | CCGTCGCGGATCCGGAAAGCTAGAAGGTTGTACAGTTAACTCCCAGGGTGTAA |
| HH-pre-miR-144 | HH-pre-144-1F | TAATACGACTCACTATAGGCTCG |
| | HH-pre-144-2R | ACCCTGATGGTGTCTGAAAAACGTACCCTGATGGTGTACGAGCCTATAGTGAGTCGTATT |
| | HH-pre-144-3F | TTCAGACACCATCAGGGTCTGTGATGATATCCCTGATGAGTCCGTG |
| | HH-pre-144-4R | CGACGGTACCGGGTACCGTTTCGTCCTCACGGACTCATCAGGGA |
| | HH-pre-144-5F | ACCCGGTACCGTCGGATATCATCATATACTGTAAGTTTGCGATGAGACACTAC |
| | HH-pre-144-6R | mAmGTACATCATCTATACTGTAGTGTCTCATCGCAAACTT |
| GG-pre-miR-19a | GG-pre-miR19a-F | TAATACGACTCACTATAGGTTTTGCATAGTTGCACTACAAG |
| | GG-pre-miR19a-R | mUmCAGTTTTGCATAGATTTGCACAACTACATTCTTCTTGTAGTGCAACTATGCAAAACC |
| Pre-miR-143 | pre-miR143-F | TAATACGACTCACTATAGGTGCAGTGCTGCATCTCTGGTCAGTT |
| | pre-miR143-R | mGmAGCTACAGTGCTTCATCTCAGACTCCCAACTGACCAGAGATGCAGCACTGCACCT |

[a] m denotes 2′-*O*-Me modification of the primer.

**Table S2. Amplification primers for template.**

| Amplification primer | 5'-sequence-3'[a] | application |
|---|---|---|
| UNIV-pUC19_E105 | TCTTCGCTATTACGCCAGCTGGCGAAA | Forward primer for amplification of DNA template for HH-pre-let-7c, HH-pre-miR-31 from plasmid |
| HDV-AMP-R | mUmAATGTGAGAATTGGCTACGTTGAAACAACGCATTACCG | Reverse primer for amplification of DNA template for HH-pre-miR-31-HDV from plasmid |
| HH-pre-let7c-AMP-R | mGmGAAAGCTAGAAGGTTGTACAGTTAACTCCCAGGG | Reverse primer for amplification of DNA template for HH-pre-let-7c from plasmid |

[a] m denotes 2′-*O*-Me modification of the primer.

**Table S3. RNA sequences.**

| RNA name | 5'-sequence-3'[a] |
|---|---|
| pre-miR-31 | AGGCAAGAUGCUGGCAUAGCUGUUGAACUGGGAACCUGCUAUGCCAACAUAUUGCCAU |
| pre-miR-21 | UAGCUUAUCAGACUGAUGUUGACUGUUGAAUCUCAUGGCAACACCAGUCGAUGGGCUGU |
| pre-let-7c | UGAGGUAGUAGGUUGUAUGGUUUAGAGUUACACCCUGGGAGUUAACUGUACAACCUUCUAGCUUUCC |
| pre-miR-144 | GGAUAUCAUCAUAUACUGUAAGUUUGCGAUGAGACACUACAGUAUAGAUGAUGUACU |
| GG-pre-miR-19a | gGUUUUGCAUAGUUGCACUACAAGAAGAAUGUAGUUGUGCAAAUCUAUGCAAAACUGA |
| pre-miR-143 | GGUGCAGUGCUGCAUCUCUGGUCAGUUGGGAGUCUGAGAUGAAGCACUGUAGCUC |

[a]Non-native residues are in lowercase.

**Table S4. Antisense oligo sequences.**

| Antisense oligo | primer names | 5'-sequence-3'[a] |
|---|---|---|
| Anti-miR31-A1 | anti-miR31-D3 | CAGGTTCCCAGTTCAACAG |
| Anti-miR31-A2 | anti-miR31-loop-D | AGGTTCCCAGTTCAACA |
| Anti-miR31-A3 | anti-miR31-D1 | GGTTCCCAGTTCAAC |
| Anti-miR31-A4 | anti-miR31-D2 | GTTCCCAGTTCAAC |
| Anti-miR31-A4-LNA | anti-miR31-D2-LNA | +G*+T*+T*C*+C*C*A*+G*T*+T*C*+A*+A*+C |
| Anti-miR144-A1 | Anti-miR144-D3 | AGTGTCTCATCGCAAACTT |
| Anti-miR144-A2 | Anti-miR144-loop-D | GTGTCTCATCGCAAACT |
| Anti-miR144-A3 | Anti-miR144-D1 | GTGTCTCATCGCAAA |
| Anti-miR144-A4 | Anti-miR144-D2 | GTCTCATCGCAAAC |
| Anti-miR144-A4-LNA | anti-miR144-D2-LNA | +G*+T*+C*T*+C*A*T*+C*G*+C*A*+A*+A*+C |
| Anti-miR19a-A2 | Anti-miR19a-loop-D | AACTACATTCTTCTTGTA |
| Anti-miR143-A4 | Anti-miR143-D2 | GACTCCCAACTGACC |
| Anti-let-7c-A1 | pre-let-7c-AP-ASO | TCCCAGGGTGTAAC |
| ASO (-) control | anti-miR-31-locked | +C*C*+G*T*T*+C*+T*A*C*+G*A*+C*C*+G*+T |
| Anti-miR-31 (+) control | Horizon Discovery | Catalog ID: IH-300507-06 |
| Anti-miR-144 (+) control | Horizon Discovery | Catalog ID: IH-300612-06 |

[a]'*' indicates phosphorothioate backbone, '+' indicates locked nucleic acid.

**Table S5. Templates for generating pmirGLO plasmid inserts.**

| Construct | Primer name | 5'-sequence-3' |
|---|---|---|
| pmirGLO miR31 | pmirGLO-miR-31-F | AAACTAGCGGCCGCTAGTAGCTATGCCAGCATCTTGCCTT |
| | pmirGLO-miR-31-R | CTAGAAGGCAAGATGCTGGCATAGCTACTAGCGGCCGCTAGTTT |
| pmirGLO miR144 | pmirGLO-miR144-N-F | GCGTTTAAACTAGCGGCCGCTAGTAGTACATCATCTATACTGTATCTAGAGC |
| | pmirGLO-miR144-N-R | GCTCTAGATACAGTATAGATGATGTACTACTAGCGGCCGCTAGTTTAAACGC |

**Table S6. Overlap extension PCR primers for generation of pCMV miR31 plasmid insert.**

| Construct | Primer name | 5'-sequence-3' |
|---|---|---|
| pCMV miR31 | cmv-miR31-1F | GCAGATCTAGTCATAGTATTCTCCTGTAACTTGGAACTG GAGAGGAGGCAAGATGCT |
| | cmv-miR31-2R | GGCATAGCAGGTTCCCAGTTCAACAGCTATGCCAGCATC TTGCCTCCTCT |
| | cmv-miR31-3F | TGGGAACCTGCTATGCCAACATATTGCCATCTTTCCTGTC TGACAGCAGCCATGGCCAC |
| | cmv-miR31-4R | GCCTCGAGGCATGCAGGTGGCCATGGCTGCTGTCAGACA GGAAA |

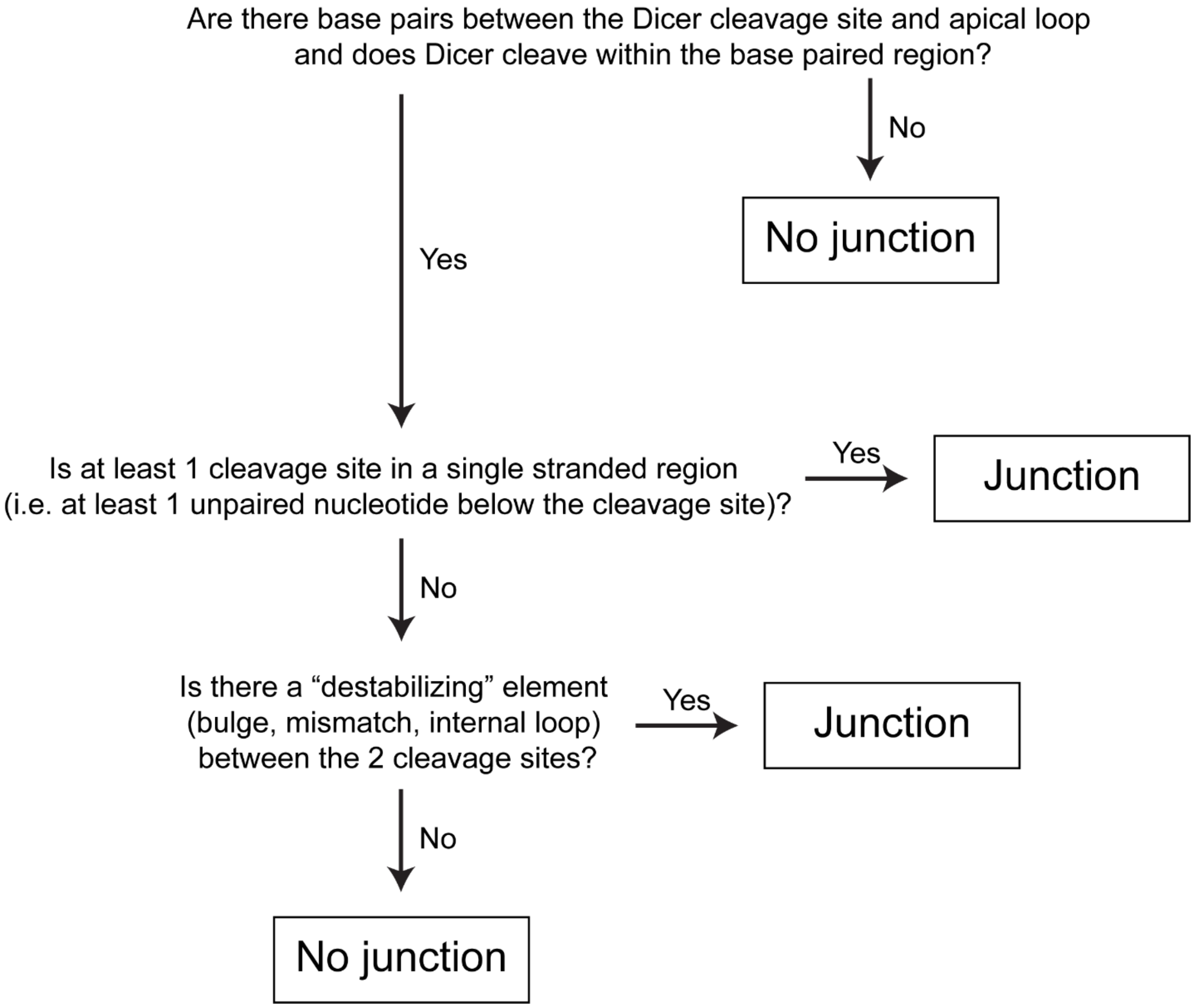


**Figure S1** Flowchart of junction region-containing pre-miRNAs determination.

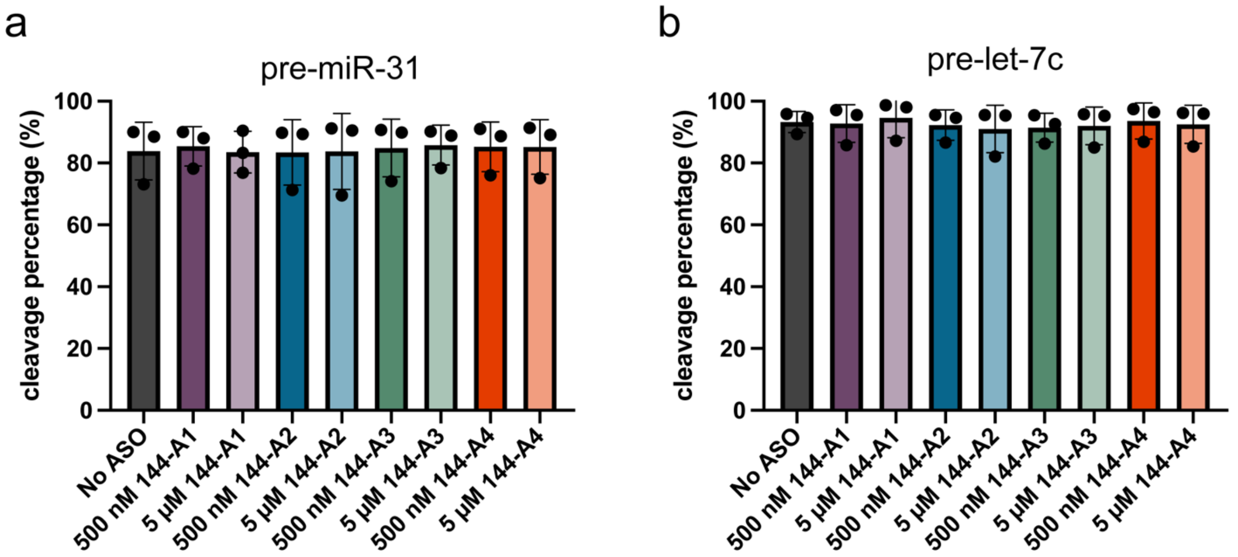


**Figure S2.** Anti-miR144 ASO are specific and do not affect the processing of pre-miR-31 or pre-let-7c. **a)** Dicer-TRBP processing of pre-miR-31 in the presence and absence of different anti-miR144 ASOs. **b)** Dicer-TRBP processing of pre-let-7c in the presence and absence of different anti-miR144 ASOs. No significant differences were identified from an ordinary one-way ANOVA Tukey analysis.

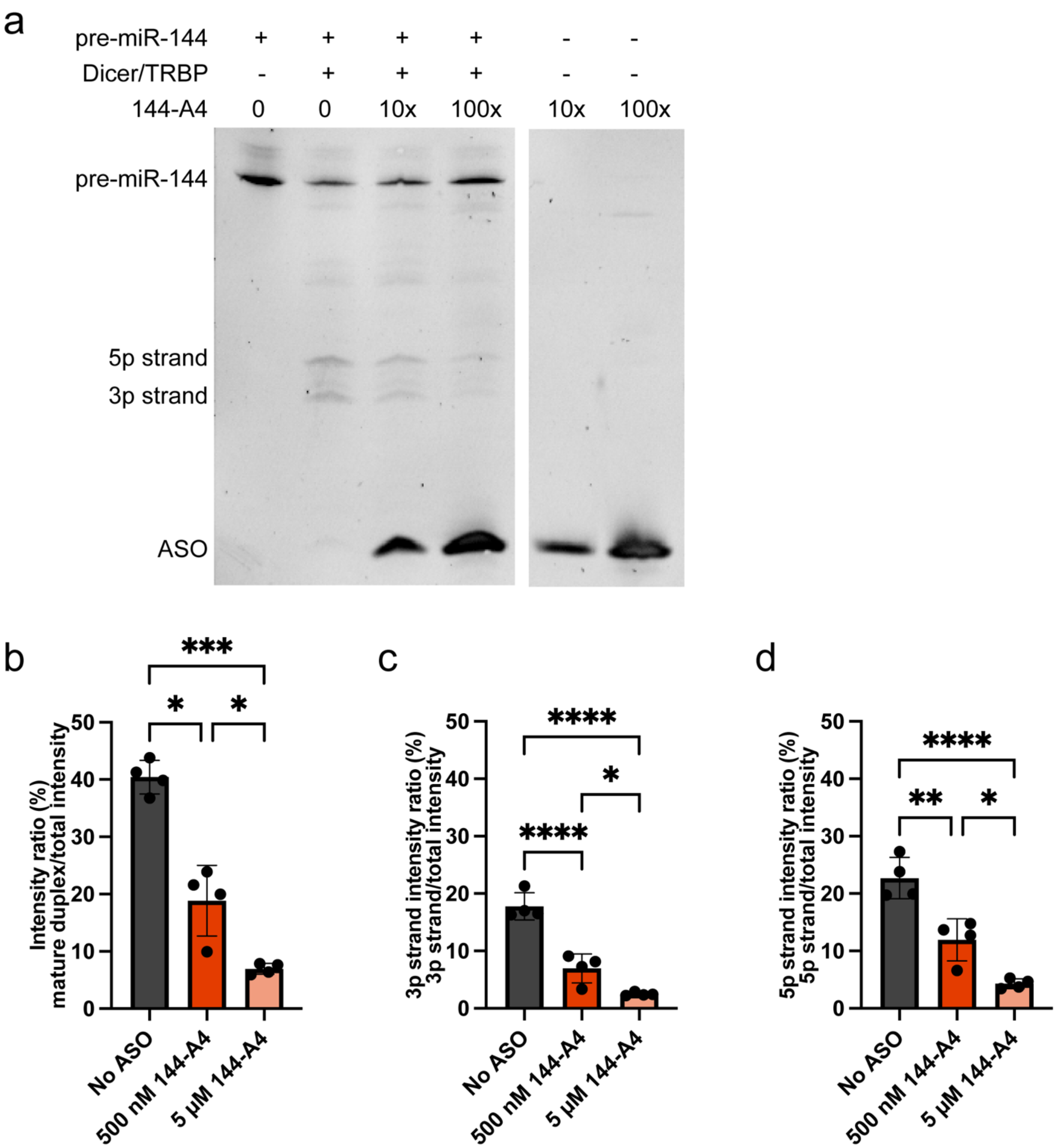


**Figure S3.** ASO 144-A4 inhibits Dicer-TRBP processing of pre-miR-144. **a)** Representative pre-miR-144 processing assay, visualized by SYBR gold staining. **b-d)** Quantification of pre-miR-144 processing products. **** $p<0.0001$, ***$p<0.001$, **$p<0.01$, and *$p<0.05$ from an ordinary one-way ANOVA Tukey analysis.

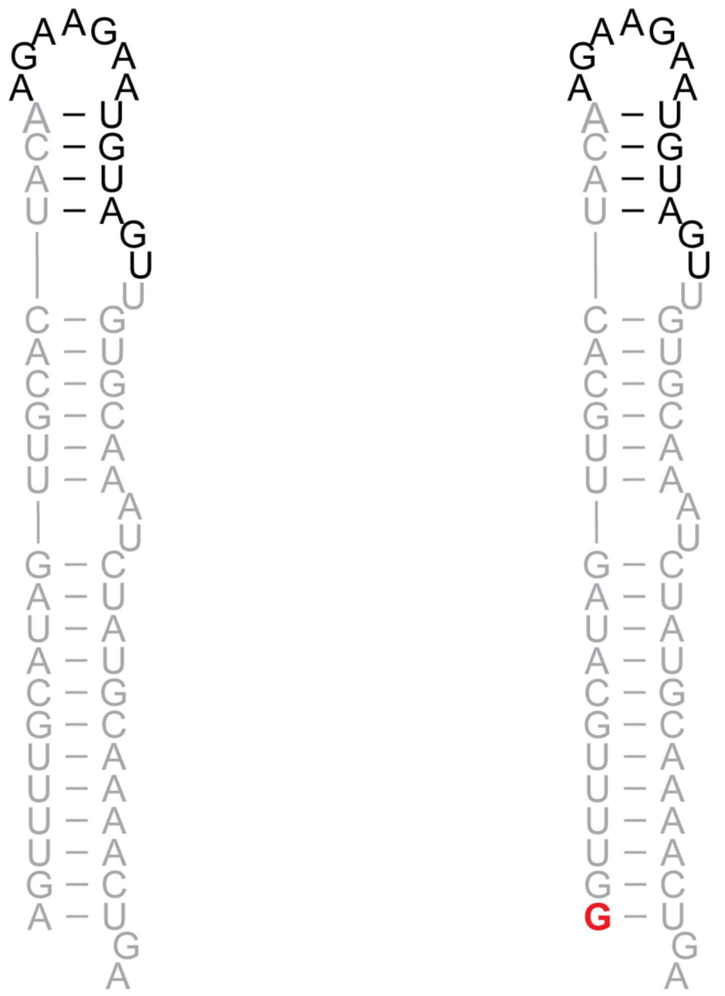

pre-miR-19a GG-pre-miR-19a

**Figure S4.** Secondary structure of pre-miR-19a and GG-pre-miR-19a.

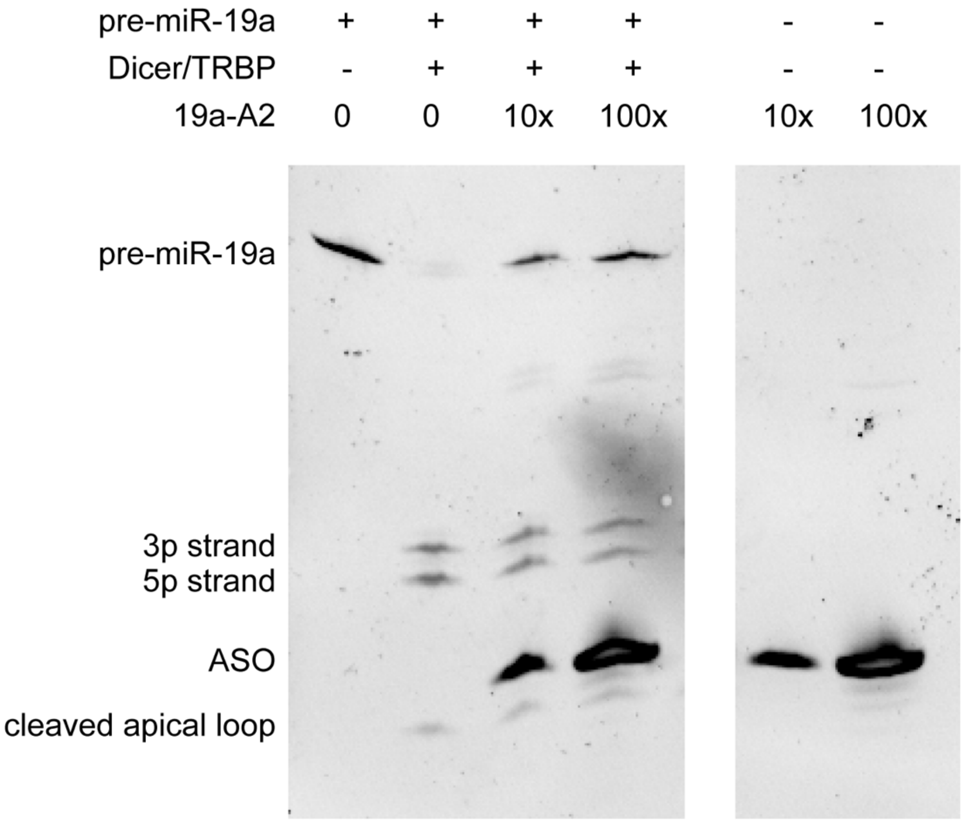


**Figure S5.** Representative GG-pre-miR-19a processing assay, visualized by SYBR gold staining.

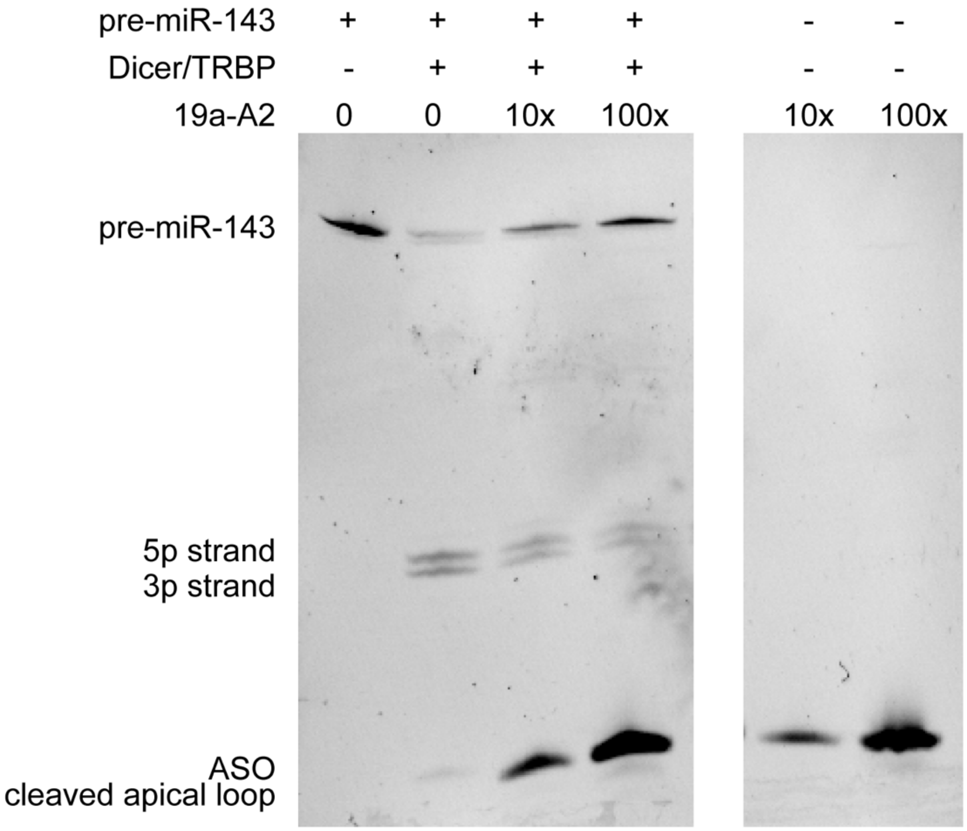


**Figure S6.** Representative pre-miR-143 processing assay, visualized by SYBR gold staining.

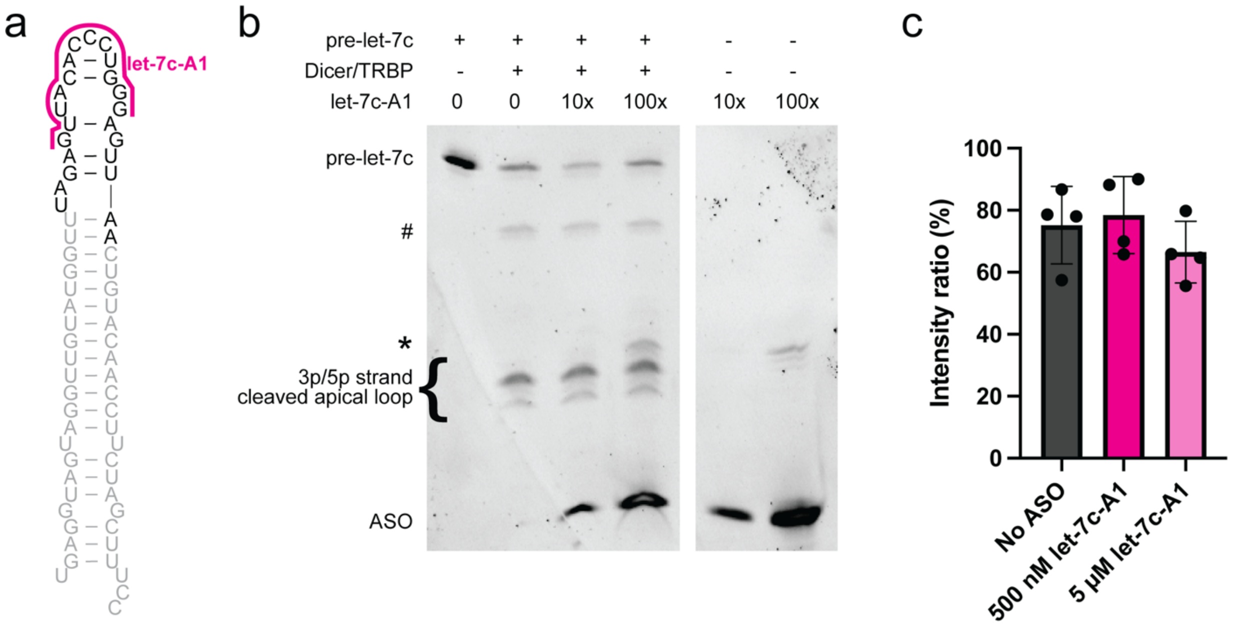


**Figure S7. a)** Predicted secondary structure of pre-let-7c. The mature let-7c duplex sequence is gray and cleaved off apical loop is black. **b**) Representative pre-let-7c processing assay, visualized by SYBR gold staining. # indicates single processed pre-let-7c product, * indicates an impurity from the anti-let-7c-A1 ASO. **c)** The intensity ratio of mature let-7c duplexes is not affected as let-7c-A1 concentration increases in the reaction.
